# Decoy-seq unlocks scalable genetic screening for regulatory small noncoding RNAs

**DOI:** 10.1101/2025.01.25.634869

**Authors:** Benedict Choi, Sushil Sobti, Larisa M. Soto, Trey Charbonneau, Aiden Sababi, Albertas Navickas, Hamed S. Najafabadi, Hani Goodarzi

**Author notes:** Correspondence: HG, and HSN. These authors contributed equally to this work (Benedict Choi, Sushil Sobti).

## Abstract

Small noncoding RNAs (smRNAs) play critical roles in regulating various cellular processes, including development, stress response, and disease pathogenesis. However, functional characterization of smRNAs remains limited by the scale and simplicity of phenotypic readouts. Recently, single-cell perturbation screening methods, which link CRISPR-mediated genetic perturbations to rich transcriptomic profiling, have emerged as foundational and scalable approaches for understanding gene functions, mapping regulatory networks, and revealing genetic interactions. However, a comparable approach for probing the regulatory consequences of smRNA perturbations is lacking. Here, we present Decoy-seq as an extension of this approach for high-content, single-cell perturbation screening of smRNAs. This method leverages U6-driven tough decoys (TuD), which form stable duplexes with their target smRNAs, for inhibition in the cell. Lentiviral-encoded TuDs are compatible with conventional single-cell RNA-sequencing (scRNA-seq) technologies, allowing joint identification of the smRNA perturbation in each cell and its associated transcriptomic profile. We applied Decoy-seq to 336 microRNAs (miRNAs) and 196 tRNA-derived fragments (tRFs) in a human breast cancer cell line, demonstrating its ability to uncover complex regulatory pathways and novel functions of these smRNAs. Notably, we show that tRFs influence mRNA polyadenylation and regulate key cancer-associated processes, such as cell cycle progression and proliferation. Therefore, Decoy-seq provides a powerful framework for exploring the functional roles of smRNAs in normal physiology and disease, and holds promise for accelerating future discoveries.

## Introduction

Small noncoding RNAs (smRNAs) represent a large family of endogenously expressed transcripts of 18-200nt that contribute to the regulation of almost all aspects of cellular biology, from developmental processes to stress response, at the transcriptional, post-transcriptional and translational levels^1^. Aberrant expressions of these transcripts are involved in many pathological conditions such as cancer, autoimmune diseases, and neurodegenerative disorders^2^. While new sequencing methods have advanced the discovery of novel smRNA transcripts in various cell and tissue types and conditions, functional characterization of these annotated smRNAs remains limited in scale to a few transcripts at a time or involves low-dimensional phenotypic readouts such as growth. There is a lack of scalable approaches that enable wider systematic mapping of the regulatory circuitry that links individual smRNAs with their related biological processes and phenotypes.

Pooled CRISPR perturbation screening coupled with single-cell RNA-sequencing has emerged as a powerful scalable approach for high-dimensional genotype-phenotype mapping in various model systems^3–6^. However, the current technology is geared towards perturbing protein-coding or long non-coding genes by suppression or activation of transcription by RNA polymerase II (Pol II). While there have been attempts to apply the CRISPR-Cas9 system to study microRNAs (miRNAs)^7^, this approach is limited only to those expressed from dedicated smRNA gene loci. Other smRNAs originate from the introns of protein-coding genes^8^ or are post-transcriptional derivatives of other RNA transcripts, including intragenic miRNAs as well as transfer RNA-derived fragments (tRFs)^9^. Current CRISPR perturbation approaches fail to uncouple perturbation effects of the derived smRNAs from those of precursor genes and are thus not suitable for high-content screens for different classes of smRNAs at large.

In the present study, we address this limitation and present a new experimental strategy for high-content single-cell perturbation screens of smRNAs termed Decoy-seq. Decoy-seq takes advantage of efficient and stable vector-encoded inhibitors—tough decoys (TuDs)^10^—to effectively inhibit the activity of smRNAs in the cell. When coupled with single-cell RNA sequencing, each perturbation can be then identified by capturing the decoy embedded in the 3’ untranslated region (UTR) of a marker gene using a targeted amplification and sequencing approach. We applied our approach to 336 miRNAs expressed in a human breast cancer cell line, showing that Decoy-seq effectively recapitulates the expected perturbation effects on the miRNA targets and reveals diverse transcriptomic phenotypes associated with different miRNA perturbations. Leveraging the power of Decoy-seq, we then generated the first high-dimensional single-cell perturbation map of 196 tRFs using the same cell line model. We derive novel insights into the diverse functional pathways governed by this poorly understood class of smRNAs, and illustrate their connections to mRNA

polyadenylation and cancer-related cellular processes such as cell cycle progression and proliferation. In sum, we present Decoy-seq as a novel, scalable platform that enables single-cell perturbation screens of smRNAs, providing a systematic and rich portrait of their regulatory functions.

## Results

### Overview of Decoy-seq design and experimental outline

The core design of Decoy-seq consists of a library of smRNA inhibitors—tough decoys (TuDs)— encoded in the 3’UTR of a Pol II marker gene (e.g., BFP) in a modified CROP-seq vector backbone^11^ where the original sgRNA scaffold is removed (Fig 1a). After transduction in target cells and selection by fluorescence-activated cell sorting (FACS), the transcriptome and perturbation identity of each individual cell are captured using a conventional polyA-based scRNA-seq technology. A TuD is a hairpin-shaped RNA decoy with a large internal bulge containing two binding sites for target smRNA^10^. We chose TuDs over other existing vector-encoded alternatives for Decoy-seq because of several advantages: 1) TuDs are experimentally tested to be the most potent inhibitor in a side-by-side comparative study^12^; 2) their inhibitory efficiency is proven in various classes of smRNAs, such as miRNAs^12^ and orphan noncoding RNAs^13^; 3) their short length is ideal for their placement in and the function of the lentiviral long terminal repeat (LTR) in the CROP-seq setup^14^; 4) their bulged structure makes them more stable and less prone to endonucleolytic cleavage^12^, and thus can achieve sustained inhibition for screening purposes; and 5) they are adaptable for high-throughput screening^15^. We reasoned that these features make TuDs suitable for perturbation screening of smRNAs, enabling high-throughput phenotypic profiling.

**Fig 1.**
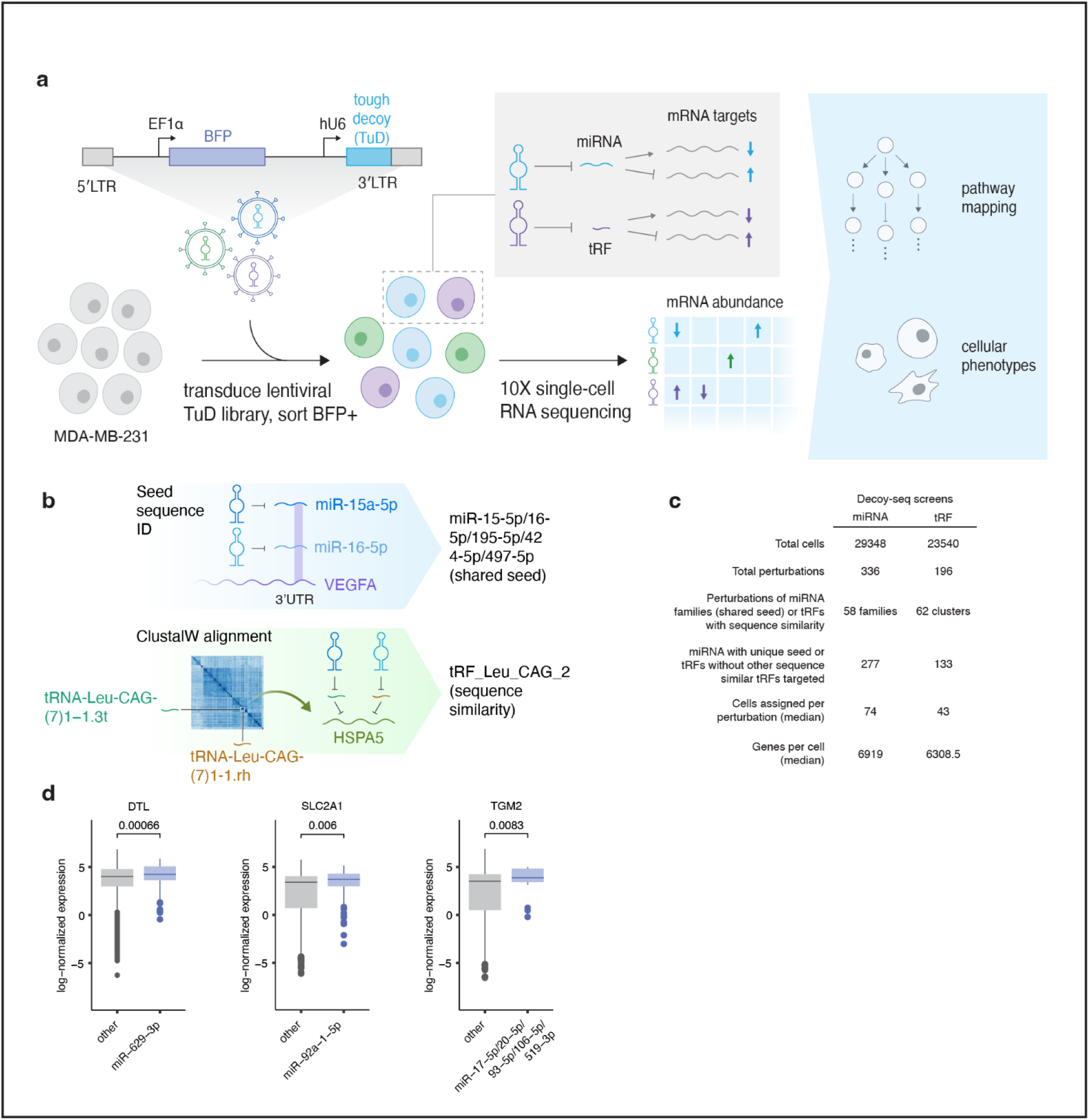
Overview of Decoy-seq. **a,** Graphical illustration of the design of Decoy-seq and experimental outline in this study. **b,** Schematic summary of the grouping of miRNAs into miRNA families based on the shared seed sequence determined by TargetScan, and the grouping of tRFs into clusters based on sequence similarity determined by ClustalW alignment. Clusters are denoted by the initial tRF in the naming whereas the initial tRNA in the naming indicates a unique non-clustered tRF. **c,** Summary information of the miRNA and tRF screens. **d,** Box-and-whisker plots of log-normalized expression of the predicted target genes (DTL, SLC2A1 and TGM2 from the left to right panel) in single cells expressing TuDs targeting other miRNAs or miR-629-3p, miR-92a-1-5p, and miR-17-5p/20-5p/93-5p/106-5p/519-3p family, respectively, in the miRNA screen. P-values calculated using two-tailed Student’s t-test.

To demonstrate the feasibility of our approach, we performed two independent screens (Fig 1a). First, a screen was performed to target miRNAs, where their functionality is well-established and their putative targets are often known. Second, we performed a discovery screen to explore an emerging class of small RNAs with poorly understood functions—tRNA-derived fragments (tRFs). These two screens targeted 336 miRNAs and 196 tRFs, respectively, expressed in the MDA-MB-231 cell line, whose smRNA expression profile we had previously determined^13^.

We profiled the pool of the cells from both screens using 10x Genomics Chromium 3’ scRNA-seq, with the addition of an enrichment PCR step for the 3’UTR region of BFP to increase the recovery of TuDs and assignment of cell perturbations^14^. The data were used to assign a unique perturbation to each single cell in each screen, along with retrieval of its transcriptome profile. Perturbation identities were defined based on miRNA “families” and tRF clusters, as opposed to individual miRNAs or tRFs, because miRNA families with the same seed sequence have largely overlapping regulons, and tRF clusters sharing similar sequences (as defined by ClustalW alignment) may be bound and inhibited by the same TuDs due to partial or imperfect base pairing (Fig 1b). After excluding low-quality cells with low UMI counts and high mitochondrial content, as well as cells bearing multiple TuDs targeting different miRNA families or tRF clusters (Extended Fig 1a-e; see **Methods**), we retained 29348 and 23540 high-quality cells covering 277 and 133 unique perturbations, with a median coverage of 74 and 43 cells per perturbation in the miRNA and tRF screens, respectively (Fig 1c).

To test whether TuDs encoded in the CROP-seq lentiviral LTR retain their function^12^, we examined the inhibitory efficacy of our library design by measuring the downstream effect on the expression of known miRNA target genes. Specifically, quantitative reverse transcription polymerase chain reaction (RT-qPCR) showed that the known target gene of miR-16-5p, VEGFA^16^, increased in expression by two folds in cells transduced with a miR-16-5p-targeting TuD encoded in the Decoy-seq setup compared to those transduced with a non-targeting (NT) control, an effect comparable to that of a commercially available TuD (Extended Fig 1f). This effect could also be observed in scRNA-seq-based data obtained from sequencing 10x Chromium libraries enriched for known targets of a select set of miRNAs (see **Methods** for experimental details). Specifically, cells with miR-16-5p perturbation expressed significantly higher VEGFA than those expressing NT controls (Extended Fig 1g), consistent with the above RT-qPCR results. Moreover, when compared to cells expressing other TuDs, we observed a significantly higher expression of DTL in cells with miR-629-3p perturbation (p-value = 0.00066, two-tailed Student’s t-test), SLC2A1 in cells with miR-92a-1-5p perturbation (p-value = 0.006, two-tailed Student’s t-test), and TGM2 in cells with perturbation of the miR-17-5p family (p-value = 0.0083, two-tailed Student’s t-test; Fig 1d), all of which represent high-confidence known miRNA-gene relationships based on TargetScan^17^. These analyses confirmed the inhibitory efficacy of our Decoy-seq libraries and the fidelity of the perturbation assignment.

### Decoy-seq captures the perturbation effects on transcriptomic phenotypes and uncovers miRNA-mediated regulatory networks

Next, we leveraged the miRNA Decoy-seq screen set data to examine the perturbation effects on transcriptomic networks at a global level (Extended Fig 2a). We aggregated the single-cell data from each miRNA perturbation and 20 pooled non-targeting controls, and generated pseudo-bulk expression profiles. We then constructed a perturbation-perturbation correlation matrix based on the pseudo-bulk expression of 3000 highly variable genes (Fig 2a; Extended Fig 2b). As expected, the transcriptomic phenotypes of perturbations of miRNAs belonging to the same family were significantly more similar than those of non-family miRNA, confirming the robustness of our screen analysis (p-value = 0.034, Wilcoxon test; Extended Fig 2c). Our analysis revealed various clusters of transcriptomic states associated with different miRNA perturbations (Fig 2a). This indicated specific miRNA perturbations could lead to distinct cellular transcriptomic phenotypes.

**Fig 2.**
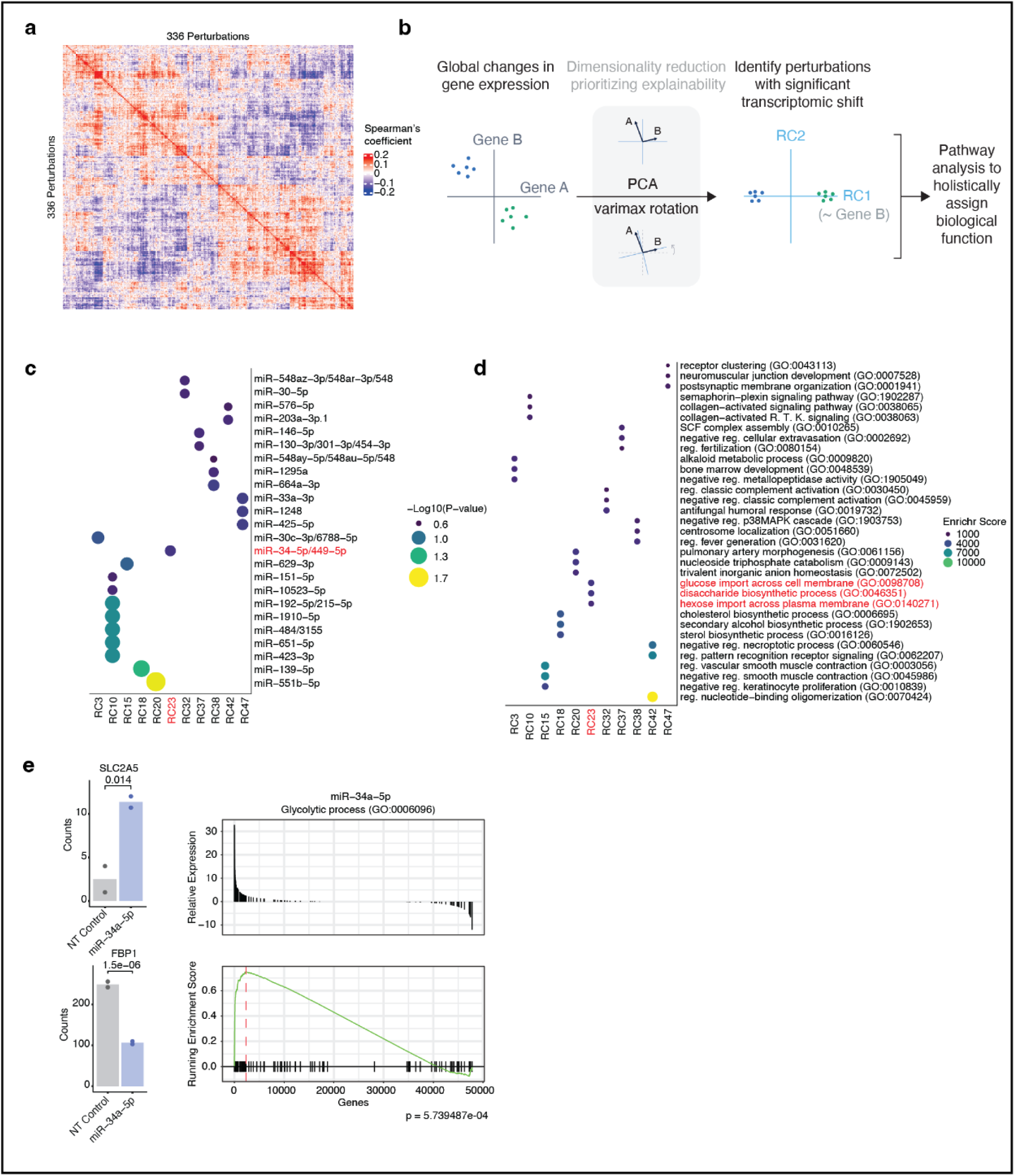
Decoy-seq delineates miRNA-mediated transcriptomic phenotypes and regulatory networks. **a,** Heatmap displaying the Spearman’s correlation of each miRNA perturbation based on the aggregated pseudo-bulk expression of 3000 highly variable genes. Each row and column represent an individual miRNA perturbation. Red and blue colors denote positive and negative correlations, respectively. **b,** Graphical illustration of the analytical framework, which incorporates principal component analysis (PCA) followed by varimax rotation to identify perturbations with significant phenotypic changes and assign the biological function. RC: “rotated component”. **c,** Dotplot showing miRNA perturbations with significant shifts in the transcriptomic states across a varimax-rotated principal component. The x-axis represents the RCs and the y-axis represents the miRNA perturbations that are best correlated to the variations in transcriptomic states captured by the respective RCs. Log-likelihood ratio test was used to obtain p-values (adjusted using Benjamini-Hochberg procedure); values passing FDR of 25% plotted. **d,** Dotplot showing a gene ontology (GO) enrichment analysis of the same RCs shown in c using Enrichr. **e,** (Left) barplots of normalized gene expression count in hexose import and glycolysis markers SLC2A5 and FBP1 from relevant gene-sets highlighted in d from the bulk RNA-seq data of MDA-MB-231 cells transduced with a non-targeting (NT) control TuD or a TuD targeting miR-34a-5p. P-values calculated with DESeq2 using the two-tailed Wald test and adjusted with the Benjamini-Hochberg procedure. (Right) Gene set enrichment analysis (GSEA) of differentially expressed genes of the glycolytic process GO geneset shown. Plotted is the relative expression or the running enrichment score of all sequenced genes ordered by upregulated to downregulated via the DESeq2 Wald test statistic. P-value calculated by odds of peak enrichment score out of GSEA simulations done with random gene order, adjusted for all gene-set comparisons by Benjamini-Hochberg procedure.

Significant changes in molecular phenotypes can take various forms, ranging from alterations in a small, targeted set of genes to broader shifts in cellular states, for example transitions between cell cycle phases. To identify significant phenotypic changes mediated by miRNAs, we performed principal component analysis (PCA) of the single-cell transcriptomes followed by varimax rotation^18^ (Fig 2b), and identified miRNA families whose perturbations were significantly associated with the rotated principal components (RCs). We identified 24 miRNA perturbations significantly associated with shifts across at least one varimax rotated component (logistic regression, FDR < 0.25, Fig 2c). The top genes with an absolute loading value greater than 0.1 from each significant component were then used for enrichment analysis to elucidate the perturbed biological pathways (Fig 2d, Extended Fig 2d; see **Methods**). Notably, several identified pathways aligned with the annotated functions of the associated miRNAs. For instance, top genes of RC37 were associated with the GO term “negative regulation of cellular extravasation”. This RC was also significantly associated with miR-146-5p and the miR-130-3p/301-3p/454-3p family, which aligned with the known roles of these miRNAs in cell migration and extravasation^19,20^. Another example was the association between top genes of RC38 and the GO term “negative regulation of p38 MAPK cascade”. Consistently, RC38 was significantly linked with miR-664-3p and the miR-548ay-5p/548au-5p/548 family, which have validated regulatory roles in the MAPK signaling^21,22^.

To further validate the robustness of our analysis, we performed bulk total RNA sequencing (RNA-seq) on the MDA-MB-231 cell line, separately transduced with TuDs specific for two of the identified significant miRNAs or a non-targeting control. Consistent with the enrichment of cholesterol biosynthesis-related pathways observed in Decoy-seq (RC18, Fig 2c-d), inhibition of miR-139-5p upregulated processes involved in cholesterol metabolism in our bulk RNA-seq data (Extended Fig 2e). For example, the expression of the gene HSD17B7, which encodes a key enzyme in the pathway, was significantly upregulated upon miR-139-5p inhibition (adjusted p-value = 0.00018; Extended Fig 2f). Similar fold expression changes were observed in the gene-set “GO cholesterol biosynthetic process” between the bulk RNA-seq and Decoy-seq data of the same miRNA inhibition (Pearson’s R = 0.37, p-value = 0.039; Spearman’s R = 0.5, p-value = 0.0051; Extended Fig 2g). Additionally, our Decoy-seq results highlighted the potential regulatory role of miR-34-5p in glycolytic pathways (RC23, GO terms “glucose/hexose import” and “disaccharide biosynthesis”; Fig 2c-d). Experimental inhibition of miR-34-5p confirmed this regulatory relationship, as demonstrated by the positive enrichment of genes involved in the glycolysis (Fig 2e, Extended Fig 2h). For example, the gene SLC2A5, which encodes the fructose transporter GLUT5 (adjusted p-value = 0.014), was upregulated while the gene FBP1, which encodes a rate-limiting enzyme in gluconeogenesis, was downregulated (adjusted p-value = 1.5×10^-6^; Fig 2e). Together, these findings support the utility of Decoy-seq in capturing smRNA-mediated regulatory networks and providing valuable insights into their biological functions.

### Revealing the multifaceted regulatory roles of tRNA-derived fragments

Having established the utility of Decoy-seq, we next focused on tRFs to uncover new biological insights. Increasing evidence suggests that tRFs play significant roles in cellular processes^9,23^, with their dysregulation linked to various diseases^24^. However, our understanding of their functional roles remains limited, and the extent of their functionality across different tRFs is still largely unknown. Our discovery perturbation screen dataset of tRFs spanning the major groups defined by their cleavage sites in their precursor forms^23^ offered a unique opportunity to investigate these interconnections on a broader scale (Extended Fig 3a).

**Fig 3.**
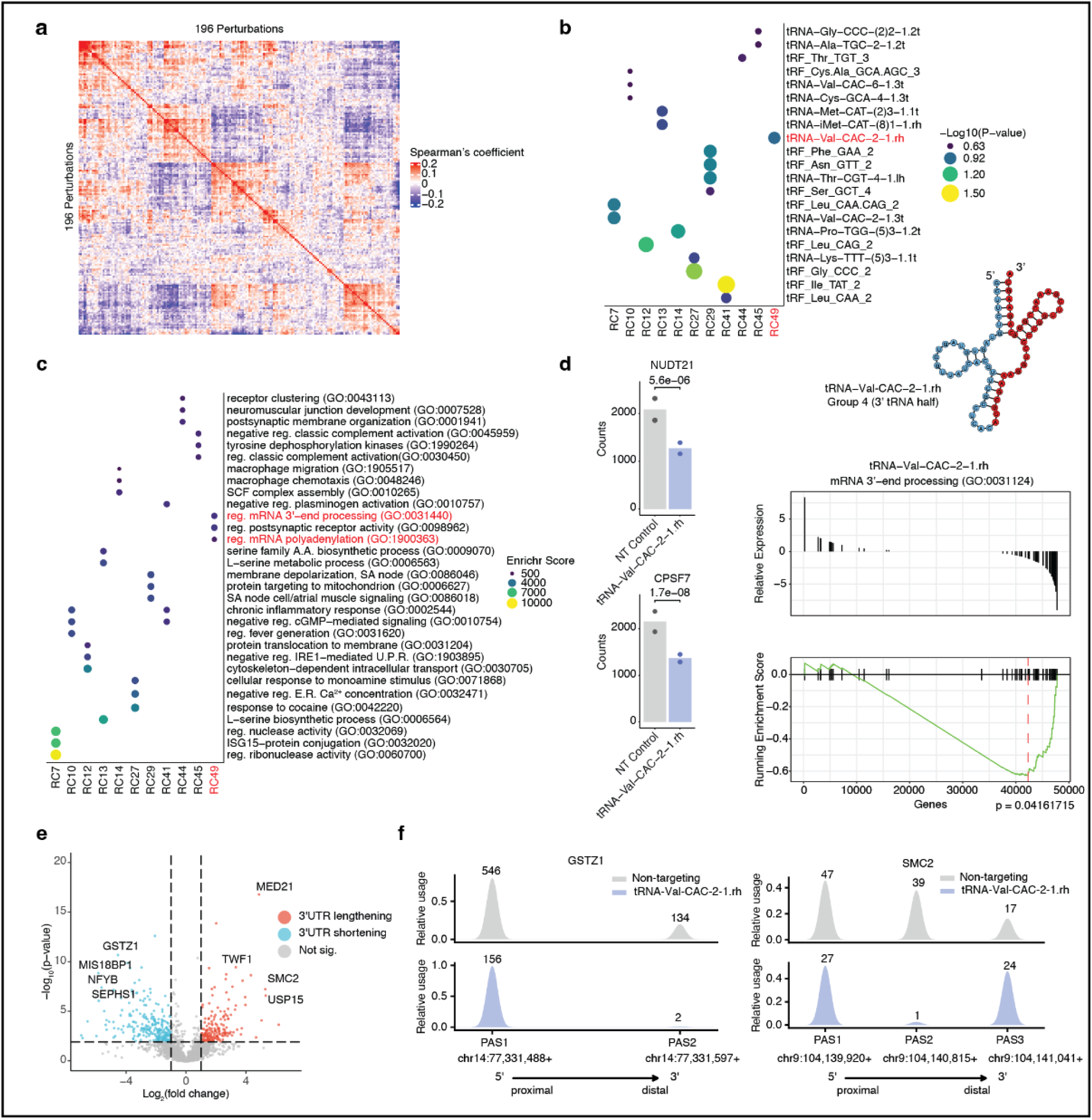
Decoy-seq reveals the multifaceted regulatory roles of tRNA-derived fragments. **a,** Heatmap displaying the Spearman’s correlation of each tRF perturbation based on the aggregated pseudo-bulk expression of 3000 highly variable genes. Each row and column represent an individual miRNA perturbation. Red and blue colors denote positive and negative correlations, respectively. **b,** Dotplot showing tRF perturbations with significant shifts in the transcriptomic states across varimax-rotated principal components. The x-axis represents the rotated components (RCs) and the y-axis represents the tRF perturbations that are best correlated to the variations in transcriptomic states captured by the respective RCs. Log-likelihood ratio test used to obtain p-values for best linear fit (adjusted using Benjamini-Hochberg procedure); values passing FDR of 25% plotted. **c,** Dotplot showing a gene ontology (GO) enrichment analysis of the same RCs shown in b using Enrichr. **d,** (Left) barplots of normalized gene expression count in mRNA 3’ processing and polyadenylation markers NUDT21 and CPSF7 from relevant gene-sets highlighted in c from the bulk RNA-seq data of MDA-MB-231 cells transduced with a non-targeting (NT) control TuD or a TuD targeting tRNA-Val-CAC-2-1.rh. P-values for change in expression calculated in DESeq2 using the two-tailed Wald test and adjusted with the Benjamini-Hochberg procedure. (Top-right) tRNA-Val-CAC-2-1 with tRF sequence tRNA-Val-CAC-2-1.rh highlighted in red and their subgrouping, indicating their origin within the tRNA from which it was derived. (Bottom-right) GSEA analysis of differentially expressed genes of the GO mRNA 3’-end processing gene-set shown. Plotted is the relative expression or the running enrichment score of all sequenced genes ordered by upregulated to downregulated via the DESeq2 Wald test statistic. P-value calculated by odds of peak enrichment score out of GSEA simulations done with random gene order, adjusted for all gene-set comparisons by Benjamini-Hochberg procedure. **e,** Volcano plot depicting differential usage of polyadenylation sites in 3’UTRs, comparing tRNA-Val-CAC-2-1.rh TuD versus non-targeting TuD pseudo-bulk cell profiles. Red dots indicate 175 3’UTR lengthening events, blue dots represent 257 3’UTR shortening events, and grey dots show non-significant changes. Statistical significance was determined with thresholds set at adjusted p-value < 0.05 and absolute log2(fold change) > 1. Key genes showing significant changes are labeled. **f,** Kernel density estimation (KDE) plots illustrating read distribution patterns across polyadenylation sites in the 3’UTR regions of two representative genes: GSTZ1 and SMC2. The x-axis shows genomic coordinates for each gene’s 3’UTR region (5’ to 3’ direction), while the y-axis represents relative usage of polyadenylation sites. Gray peaks represent non-targeting TuD cells, while blue peaks show tRNA-Val-CAC-2-1.rh TuD cells. Numbers above peaks indicate the number of reads assigned to each polyadenylation site (PAS) by MAAPER.

The perturbation-perturbation correlation matrix for each tRF revealed distinct clustering patterns, showing clear organization of various changes in transcriptomic phenotypes (Fig 3a, Extended Fig 3b). As expected, perturbation of tRFs in the same clusters (Fig 1b) produced more similar transcriptomic phenotypes than tRFs of different clusters, further confirming the robustness of our results (p=0.015, Extended Fig 3c). As described above, we performed varimax rotation analysis to identify the main axes of variation in our data and their associated tRFs and pathways. This analysis revealed 11 RCs associated with 21 unique tRF clusters, with a wide array of biological processes, ranging from RNA processing to inflammatory responses, enriched among the top genes of these RCs (Fig 3b-c, Extended Fig 3d). Many of the identified processes have not been previously linked with tRFs.

One notable example was the novel connection between tRNA-Val-CAC-2-1.rh and mRNA 3’end polyadenylation (through RC49, Fig 3b-c, Extended Fig 3d-f). To explore this relationship further, we inhibited tRNA-Val-CAC-2-1.rh in MDA-MB-231 cells and profiled the gene expression changes using bulk total RNA-seq. In line with the Decoy-seq results, inhibition of tRNA-Val-CAC-2-1.rh led to expression changes in genes enriched for mRNA 3’end processing (Fig 3d, Extended Fig 3e). Among them, the expression of the core cleavage and polyadenylation factors, including NUDT21 and CPSF7, significantly decreased (adjusted p-values = 5.6×10^-6^ and 1.7×10^-8^, Fig 3d). Because the scRNA-seq assay we used was a 3’tag-based method that generated reads near the polyadenylation sites, this allowed us to study the alternative polyadenylation (APA) changes associated with tRNA-Val-CAC-2-1.rh inhibition. Accordingly, using the aggregated pseudo-bulk data, we found that tRNA-Val-CAC-2-1.rh inhibition significantly altered the APA site usage across the transcriptome, with many transcripts undergoing 3’UTR shortening and lengthening (257 and 175 genes, respectively; Fig 3e). For instance, the distal polyadenylation site (PAS) of the gene GSTZ1 showed a significant reduction in usage, while the opposite trend was observed for SMC2 (Fig. 3f). These results underscore the regulatory role of tRNA-Val-CAC-2-1.rh in APA dynamics. In addition, we confirmed the involvement of the tRF-Leu-CAG-2 cluster in the regulation of IRE1-mediated unfolded protein response (RC12, Fig 3b-c, Extended Fig 3f-h): RNA-seq analysis revealed that inhibition of tRNA-Leu-CAG-(7)1-1.3t, a member of this cluster, in MDA-MB-231 cells led to the enrichment of gene-sets associated with the GO terms “protein folding” (p-value = 0.004993511) and “protein folding in ER” (p-value = 0.011117331, Extended Fig 3g), respectively. This was exemplified by an increase in the expression of DNAJB9 and HSPA5, both of which are known repressors of ERN1/IRE1^25,26^ (Extended Fig 3h). Collectively, our results provided compelling evidence for the complex regulatory roles of tRFs in maintaining cellular homeostasis, and the ability of Decoy-seq to reveal these molecular phenotypes.

### tRNA-derived fragments are modulators of cell cycle progression and proliferation

Previous studies have highlighted a connection between tRFs and cancer growth and progression (reviewed in ref ^27^). We expanded on these findings, leveraging our Decoy-seq dataset to explore this relationship more systematically and provide a more complete picture of the coordinated modulatory effects of tRFs. Specifically, we focused on the hallmark gene-sets^28^, which encompass major pathways frequently dysregulated in cancer, and generated an expression matrix for each tRF perturbation based on the aggregated pseudo-bulk profiles (Fig 4). This analysis revealed that many of the tRFs examined in our Decoy-seq experiment may act as modulators of one or more of these hallmark pathways.

**Fig 4.**
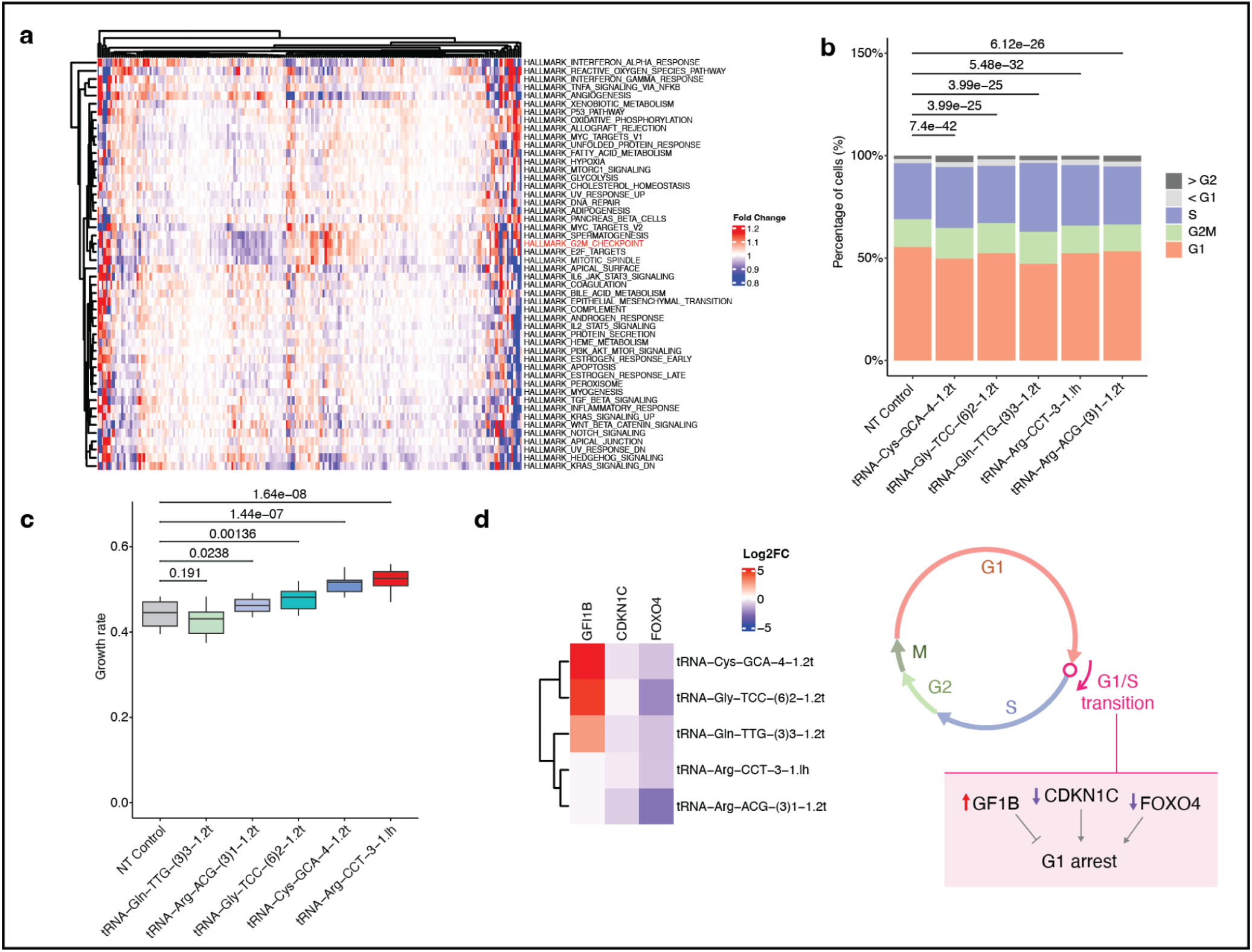
tRFs regulate cell cycle progression. **a,** Pseudobulk heatmap of mean fold change in gene expression of hallmark pathways (y-axis) across all tested tRF perturbation (x-axis). The intensity of red and blue colors denote the fold change. **b,** Relative proportion of MDA-MB-231 cells in each cell cycle phase obtained using flow cytometry of cells transduced with a targeting or non-targeting TuD. P-value calculated using chi-square test to compare homogeneity of proportions of cell cycle phases and adjusted with the Benjamini-Hochberg procedure. **c,** Box-and-whisker plot showing the growth rate of replicates of MDA-MB-231 cells transduced with a tRF targeting or non-targeting TuD. P-value calculated using two-tailed Student’s t-test and adjusted with the Benjamini-Hochberg procedure. **d,** (Left) Heatmap of differential expression of selected effector genes regulating G1 cell phase from bulk RNA-seq data of MDA-MB-231 cells transduced with respective tRF-targeting TuD versus non-targeting control. Red and blue colors denote positive and negative log2 fold changes, respectively. Generally, GF1B inhibits G1 arrest while CDKN1C and FOXO4 promote it. Together, upregulated GF1B and/or downregulated CDKN1C and FOXO4 indicate depletion of cells in G1. (Right) Illustration of perturbed genes accelerating G1/S transition.

One illustrative case was the differential expression of gene-sets associated with cell cycle progression, including the G2/M checkpoint, E2F targets, and the mitotic spindle, across various tRFs (Fig 4). We focused on tRFs that ranked within the top or bottom 10% based on their median cell cycle scores in the S or G2/M phases for experimental validation in MDA-MB-231 cells (Extended Fig 4a). Cell cycle analysis by flow cytometry demonstrated that inhibition of each of these five nominated tRFs promoted progression from G1-to-S and G2/M phases compared to the non-targeting control (Fig 4b, Extended Fig 4b-c). In line with this, inhibition of these tRFs, except for tRNA-Gln-TTG-(3)3-1.2t, significantly increased cell proliferation (Fig 4c). To uncover the downstream effector genes, we transduced MDA-MB-231 cells with a non-targeting control or targeting TuD specific for one of these five tRFs and performed bulk total RNA-seq. We compared the bulk gene expression profiles between tRF inhibition and control conditions. This analysis revealed that the observed enhanced G1/S transition and increased proliferation were driven by the upregulation of the positive regulator GFI1B^29^, and/or downregulation of the negative regulators CDKN1C^30^ and FOXO4^31^ (Fig 4d). Notably, while these validated tRFs originated from different tRNA decoders and had distinct sequences (i.e., they did not belong to the same tRF cluster), the majority of them were classified into the same subgroup (Group 6 internal tRF^19^) and shared a similar secondary structure (Extended Fig 4b). This common structural feature may be biologically important for their regulatory functions.

Taken together, our results confirmed the repressive roles of the tested tRFs in cell cycle progression through a mechanism distinct from previously reported pathways^32^, and in contrast to the positive modulatory roles described elsewhere^33^. Thus, our Decoy-seq provided valuable insights into the complex regulatory relationship between tRFs and cancer-associated phenotypes.

## Discussion

Understanding gene expression, their interactions and regulation, and how these factors influence the phenotype of a given cell is a fundamental goal in genetics and requires moving beyond traditional linear models of gene function. Perturb-seq and its variants, including CROP-seq and Direct-capture-seq^34^, are transformative tools in accelerating systematic studies of these relationships, and the release of the genome-wide Perturb-seq data^6^ marks a significant milestone in these efforts. In this study, we extended the capabilities of Perturb-seq to include small noncoding RNAs - a crucial node in (post-)transcriptional regulation - by introducing a novel approach called Decoy-seq. This innovative method facilitates a more comprehensive mapping of noncoding genome-phenotype interactions and the construction of these complex non-linear relationships into manifolds^35^ that illuminate all possible “destinies” or states of a given cell. Unlike existing CRISPR-Cas9-based techniques, which perturb transcription, Decoy-seq directly inhibits the interaction between smRNA transcripts and their cognate interactome through base pairing and stable duplex formation. This distinct mechanism makes Decoy-seq broadly applicable to all classes of smRNAs, particularly those that are processed post-transcriptionally from precursor transcripts with distinct functions. Additionally, Decoy-seq is highly flexible, compatible with any polyA-capturing single-cell sequencing technology (e.g., microfluidics and split-pooling), further enhancing its utility in high-throughput studies.

We demonstrated the utility of using Decoy-seq to capture the broad spectrum of transcriptomic phenotypes regulated by two distinct classes of smRNAs. By analyzing these phenotypes, we were able to identify their regulons, recapitulating known biological mechanisms and uncovering novel insights. Specifically, we focused on tRFs, and our data provided a compelling snapshot of their complex regulatory functions in diverse fundamental biological processes, including a novel role in mRNA polyadenylation. Importantly, our data enabled an all-in-one multi-dimensional mapping of all of these interactions in the given cell model system. This gave us a unique opportunity to explore the concerted redundant functions of different tRFs in controlling cancer-related processes, such as cell cycle progression and proliferation. Collectively, these examples suggest that the biological impact of tRFs may be far greater than previously recognized. We envision that our method will accelerate future discoveries, providing a clearer understanding of their roles in normal physiology and disease.

There has been growing interest in using machine learning models to enhance our understanding of gene regulation and predict perturbation effects using multi-modal omics data. Our method offers a scalable platform to assess different classes of regulatory small RNAs across various cell, tissue, and organ models, providing an excellent opportunity to address a significant gap in our understanding of gene regulatory circuitry. As with other Perturb-seq experiments, a major limitation is the cost associated with increasing scale. However, analytic approaches like compressed Perturb-seq^36^, combined with advances in single-cell sequencing, hold great promise for mitigating this issue and enhancing experimental scalability.

In summary, our study presents a robust framework for the systematic exploration and analysis of small noncoding RNAs and their cellular functions, contributing to a comprehensive understanding of gene regulation.

## Experimental Model and Subject Details

### Cell culture

All cells were cultured in a 37 °C 5% CO_2_ humidified incubator. The HEK293T cell line (ATCC CRL-3216) and MDA-MB-231 (ATCC HTB-26) cultured in DMEM high-glucose medium supplemented with 10% fetal bovine serum (FBS), glucose (4.5 g L^-1^), L-glutamine (4 mM), sodium pyruvate (1 mM), penicillin (100 units mL^-1^), streptomycin (100 μg mL^-1^) and amphotericin B (1 μg mL^-1^) (Gibco). All cell lines were routinely screened for mycoplasma with a PCR-based assay and tested negative.

### Cloning

The mU6-gRNA cassette of the CROP-seq vector pBA950 (Addgene #122239) was replaced by a hU6 promoter followed by AgeI and EcoRI cloning sites. Tough decoys were synthesized either as individual single strand oligos or oligo pools (Integrated DNA Technologies or Twist Bioscience) with 5’-TCTTGTGGAAAGGACGAAACACCGGT and 5’-GAATTCTCGACCTCGAGACAAATGG overhangs and Gibson-cloned into the modified CROP-seq vector digested with AgeI and EcoRI. The Gibson assembly product was purified, transformed and amplified. For Perturb-seq, the purified pooled libraries were verified on a Miseq run at UCSF Center For Advanced Technology (CAT) prior to screening. The tough decoy sequences used in this study are provided in Supplementary Table 1.

### Lentiviral production and titration

The transfer plasmid was co-transfected with plasmids pMD.2G and pCMV-dR8.91 using TransIT-Lenti (Mirus) into HEK293T cells according to the manufacturer’s manual. Viral supernatant was collected 48 hours after transfection and passed through a 0.45 µm filter. For viral titration of the perturb-seq libraries, target cells were transduced in a 6-well plate with varying amounts of viral supernatant overnight in the presence of 8 µg mL^−1^ polybrene (Millipore) and checked 48 hours post-transduction for the proportion of BFP+ cells through flow cytometry.

### Perturb-seq screen

MDA-MB-231 cells were transduced at a multiplicity of infection (MOI) of ∼0.2. After 48 hours post-transduction, BFP+ cells were isolated through fluorescence-activated cell sorting (FACS) and used directly for scRNA-seq with the 10x Genomics Chromium Next GEM Single Cell 3’ HT assay v3.1. The gene expression libraries were prepared according to the manufacturer’s manual and sequenced on a NovaSeq 6000 instrument at Chan-Zuckerberg Biohub. To assign tough decoys to single cell transcriptomes, a series of three successive PCR reactions were performed on the cDNA from the scRNA-seq assay to enrich for the tough decoy sequence in the 3’UTR of BFP transcripts as previously described^14^. The enrichment libraries were sequenced on a NextSeq 500 instrument at UCSF.

### Targeted enrichment of scRNA-seq libraries

Predicted conserved targets (mRNA) for miRNAs were obtained from TargetScan 6.0, and a panel of hybrid capture probes against miRNA target genes were designed and purchased from Twist Bioscience. The enrichment libraries were prepared using the cDNA from the 10x Genomics Chromium Next GEM Single Cell 3’ HT assay v3.1 as input according to the Twist target enrichment standard hybridization v2 protocol. The final libraries were sequenced on a NextSeq instrument at UCSF.

### Inhibition of small RNA

MDA-MB-231 cells were incubated with viral supernatant overnight in the presence of 8 µg mL^−1^ polybrene (Millipore). After 48 hours post-transduction, BFP+ cells were FACS-sorted for RNA isolation or subculturing. For the RT-qPCR experiment, a commercially available tough decoy against miR-16 (Sigma-Aldrich) was used to transduce MDA-MB-231 cells and selected with 1.5 µg mL^−1^ puromycin (Gibco).

### RNA isolation

Total RNA for RT–qPCR and RNA-seq was isolated using the Zymo QuickRNA isolation kit with in-column DNase treatment per the manufacturer’s protocol.

### Bulk total RNA-seq

RNA-seq libraries were prepared using the SMARTer Stranded Total RNA-seq kit v3 (TaKaRa Bio) following the manufacturer’s instructions. The libraries were sequenced on a NovaSeqX instrument at UCSF CAT.

### RT–qPCR

Transcript levels were measured using RT–qPCR by reverse transcribing total RNA to complementary DNA (Maxima H Minus RT, Thermo), then using LightCycler 480 SYBR Green I Master (Roche) per the manufacturer’s instructions. HPRT1 was used as the endogenous control. The primer sequences are as follows.

HPRT1_F: CCTGACCAAGGAAAGCAAAG

HPRT_R: GACCAGTCAACAGGGGACAT VEGFA_F: GCCAGCACATAGGAGAGATG VEGFA_R: CCCCTTTCCCTTTCCTCGAAC

### Cell proliferation assay

Cell proliferation was determined by high-content imaging using the IncuCyte SX5 instrument (Satorius) according to the manufacturer’s instructions. In all experiments, cells were seeded into a 96-well culture plate and for each well four fields were imaged under ×10 magnification every 6 hours. The IncuCyte software (v2020C) was used to calculate the confluency values and the cell growth rate was normalized to the cell confluency at time 0.

### Cell cycle analysis

Cells were stained with Vybrant DyeCycle Green stain (ThermoFisher) according to the manufacturer’s instructions. In brief, cells were resuspended in complete medium at a concentration of 1×10^6^ mL^-1^ and incubated with the dye at a final concentration of 10 μM at 37 °C for 30 minutes in the dark. Stained cells were analyzed on a flow cytometer and gated according to unstained samples. The FlowJo (BD Biosciences) software was used to perform the cell cycle analysis.

## Quantification and Statistical Analysis

### Alignment, cell calling, and guide assignment

Cell Ranger 7.0.0 software (10x Genomics) was used in the alignment of scRNA-seq reads to the transcriptome, alignment of TuD reads to the library, collapsing reads to UMI counts, and cell calling. The 10x Genomics GRCh38 version 2020-A genome build was used as the reference transcriptome. Geomux was used to assign TuDs to cells as shown in Extended Fig 1a.

Downstream analyses were performed in Python and R, using a combination of numpy, pandas, scanpy, Seurat, tidyverse, ggplot2, ggpubr, ggrepel, gridExtra, circlize, ggtree, cowplot, aplot, patchwork, stringr, igraph, rstatix, EnhancedVolcano, ggbreak, ggforce, escape, enrichR, SummarizedExperiment, GSEABase, DESeq2, tximport, and clusterProfiler.

### Grouping of miRNA families and tRF clusters

For groupings, TargetScan 6.0^17^ miR family annotations were used to classify miRNAs into families and the ClustalW algorithm^37^ (with default parameters) was used to cluster tRFs with high sequence similarity (> 97%) as shown in Fig 1b. The lists of miRNA families and tRF clusters and their members are indicated in Supplementary Table 2.

### miRNA target gene expression

Using data from the scRNA-seq libraries that were enriched for conserved targets of miRNAs, we tested for the differential expression of each mRNA in cells in which the targeting miRNA was perturbed, in comparison to all other cells. For this, we first used GEDI^39^ (in the ‘Bl2’ mode, with the number of latent factors ‘K’ set to 50) to normalize mRNA levels in each cell and denoise the data, resulting in a denoised measure of mRNA abundance in logarithmic scale. We used a two-sided t-test to compare the log-normalized expression values between two cell groups, followed by FDR correction using the Benjamini–Hochberg procedure. Significant differences at FDR<0.2 are reported.

### VEGFA gene expression

Seurat’s SCTransformed expression values were used to assess changes in VEGFA expression. Total number of cells recovered for miR-16-5p TuD (20) were sampled from non-targeting TuD cells to control for cell number differences. SCTransform normalized values were plotted and one-tailed Wilcoxon rank-sum test used to compute the p-value of miR-16-5p TuD median cell VEGFA expression being greater than that of non-targeting TuD.

## Decoy-seq analysis

### Differential expression and perturbation-perturbation correlation

After alignment, cell calling, and guide assignment, expression counts were normalized using Seurat’s SCTransform function. Gene expression differences across perturbations were examined using pseudo-bulked normalized counts across top 3000 variable genes in the variable genes heatmaps for miRNA and tRF perturbations. Perturbation-perturbation correlation heatmaps were constructed for miRNA and tRF perturbations by correlating each perturbation’s normalized gene expression across the 3000 variable genes with all other perturbations.

### Varimax rotation analysis

Normalized counts for top variable genes were used as input into PCA analysis to put cells into principal component space (from the original gene expression space), with 50 components being retained. Loadings of each principal component axis were then rotated using the varimax rotation to maximize correlation of loading genes to each principal component in order to improve interpretability of gene expression shifts caused by perturbed cells moving across a component axis. Rotated loadings were then used to transform the cell coordinates in principal component space to rotated component coordinates in the new space. Logit models (logistic regression) were built correlating perturbation cells vs the non-targeting control cells to a given rotated component. This was repeated for all 50 rotated components. Perturbations with significant correlation to a component via p-values from the log-likelihood ratio test (FDR > 25%) were plotted in Fig 2c and 3b dotplots. For each significant perturbation, genes of the correlated component with absolute loading values > 0.1 (‘signal genes’) were inputted into Enrichr to identify gene sets (pathways) associated with them. The top 3 pathways by Enrichr score associated with the ‘signal’ genes from a perturbation were retained and plotted in the Fig 2d and 3c dotplots. The combined score (Enrichr score) is calculated in the enrichR package by taking the natural logarithm of the p-value of the Fisher exact test and multiplying it by the z-score of the deviation from the expected rank of the enriched genes in the ‘signal’ gene list.

### miRNA Decoy-seq variational autoencoder UMAP

To map the breadth of transcriptomic states accessed by miRNA perturbations, a variational autoencoder neural network was trained with expression data and perturbation identity of individual cells in the miRNA Decoy-seq normalized counts matrix. The variational autoencoder in the CiberATAC^38^ package was used as the architecture of the model with the mRNA target gene set for each miRNA family in TargetScan 6.0 annotations used to connect the input layer (genes) to the second layer (genesets). The center layer of the encoder contained 10 nodes and the dimensionality reduced data in these nodes from the training dataset was extracted and put into the UMAP algorithm to further reduce the dimensionality to 2 dimensions (Extended Fig 2a).

### Bulk total RNA-seq analysis

For bulk RNA-seq follow up of select miRNA and tRF TuDs, the Salmon mapping tool was used in quasi-mapping mode to index and quantify fastq reads with default parameters using human Gencode v33 (GRCh38.13) transcript sequences. DESeq2 was used to calculate log2 fold changes in gene expression from the quantified reads between non-targeting control and tRF targeting TuD cells. GSEA analysis (implemented via the clusterProfiler package) was then used to evaluate significant pathway gene expression changes.

### Intra-family and intra-cluster expression profile correlation

Spearman’s correlation coefficients were obtained from pseudo-bulked gene expression profiles (Seurat’s SCTransformed values used to pseudo-bulk); profiles of TuD against miRNAs in a given family were correlated to each other in all possible combinations. Median Spearman’s coefficients were obtained for each family. These were compared to median coefficients from randomly selected unique miRNA TuDs distributed into equivalently sized miRNA groups to serve as a control. This process was also repeated for TuDs against tRFs within a sequence cluster vs those in unique tRFs.

### Correlation between Decoy-seq and bulk RNA-seq data

Cells with miR-134-5p and tRF-Val-CAC-2-1.rh TuD were compared to cells with non-targeting TuD for genes in the cholesterol biosynthetic process and mRNA 3’ end processing pathway to obtain fold change values, respectively. Values for Decoy-seq and bulk RNA-seq experiments obtained from Seurat’s ‘Fold Change’ function and DESeq2, respectively. A simple linear model was fitted using ggpubr’s stat_cor function to obtain Pearson’s and Spearman’s R and p-value.

### tRF groupings and visualization of tested tRFs on their tRNAs

Groupings of tRF sequences targeted were made by tDRnamer^40^ based on the Sprinzl base pair alignment position of the tRF within its originating tRNA. Illustrations of tested tRFs, highlighted within the overall tRNA, were also obtained from tDRnamer. tRNA-Val-CAC-2-1.rh illustration in Fig 3 was made using the RiboSketch software^41^.

### Alternative polyadenylation analysis

Pseudo-bulk alignment files were generated for each condition using the cell barcodes. We employed the MAAPER software^42^ to assign sequencing reads to known polyA sites (PAs), as defined in the PolyA DB v3 database^43^. PAs sites were considered only if there were at least 25 reads aligning to the sites. To identify genes with significant changes in the length of their 3’-most exon, we used the REDu metric provided by MAAPER. REDu measures the log2 fold change or relative expression levels between the two most differentially expressed isoforms in the 3’-most exon. A positive REDu value indicates transcript lengthening events, while a negative value points to shortening events.

The RED score, comparing conditions 1 and 2, is computed using the formula:

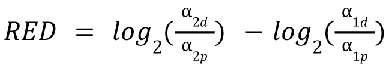

where the proportions of the distal and proximal PAs in conditions 1 and 2 are denoted as α_2𝑑_, α_2𝑝_, α_1𝑑_, and α_1𝑝_ respectively.

### Hallmark pathway expression

Seurat normalized values were used for gene expression to pseudo-bulk Decoy-seq cells by TuD identity. Genes in the GSEA’s hallmark gene set collection were kept and averaged within their respective geneset to create pathway expression. Pathway expression values for each TuD are divided by those of non-targeting TuD; values are displayed in log2 fold change format.

### Evaluation of cell cycle shifts in Decoy-seq

Cells in the dataset had S and G2M phase scores calculated using Seurat’s CellCycleScoring function on normalized counts. For each TuD, 10 random cell samplings (n = 30) were performed for all TuDs with greater than 30 cells recovered using 10 seeds for reproducibility. The median S or G2M phase score was calculated for each TuD from the 10 samplings. To create a background (‘noise floor’) set of cell samplings, cells for each TuD were again sampled (n = 30) 10 times using the same seeds, however this time cells were sampled from all TuDs in the dataset creating TuD samplings of random identity cells. These two sets of samplings were then combined. For each sampling seed, TuDs that fell in the top 10th or bottom 10th percentile were kept and the frequency of TuDs to appear in the top 10th or bottom 10th percentile was calculated across the 10 sampling seeds (Extended Fig 4a).

Cell cycle gene UMAP created from S and G2M phase scores run through Seurat’s RunUMAP function (Extended Fig 4c).

## Data availability

All sequencing data have been deposited in the GEO database under accession GSEXXXXXX.

---

**Extended Fig 1.**
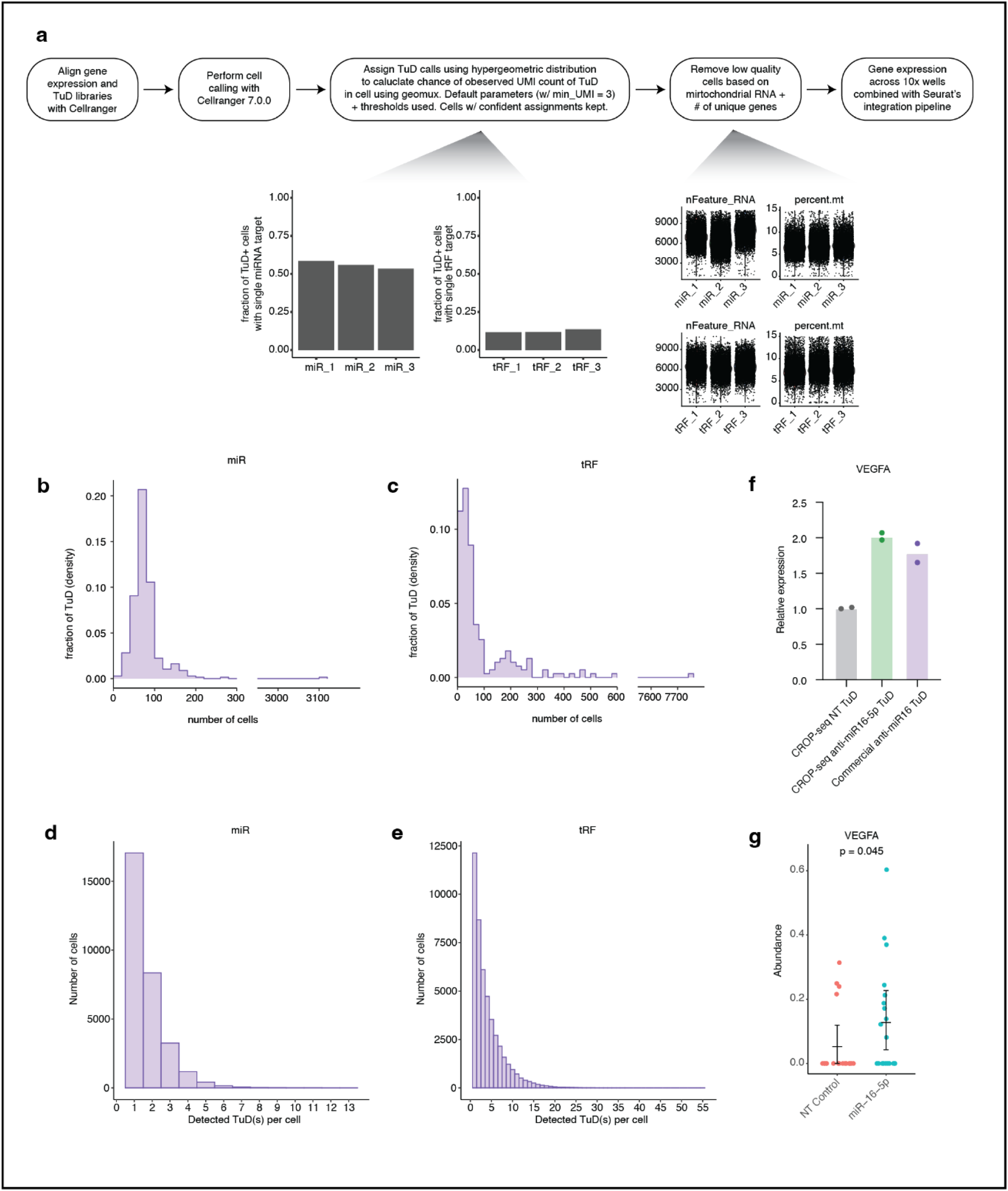
Processing and quality control of Decoy-seq screen data. **a,** Schematic overview of data processing from data alignment, cell calling, TuD assignment to filtering. **b,** Histogram showing the number of cells assigned in the miRNA Decoy-seq screen for cells expressing a given TuD; TuDs targeting miRNAs with the same seed sequence (family) are pooled together as one perturbation. **c,** Histogram showing the number of cells assigned in tRF Decoy-seq screen for cells expressing a given TuD; TuDs with high sequence similarity (cluster) determined by the ClustalW algorithm are pooled together as one perturbation. **d,** Histogram showing the distribution of the number of detected TuD(s) per cell in the miRNA screen dataset. **e,** Histogram showing the distribution of the number of detected TuD(s) per cell in the tRF screen dataset. **f,** Relative expression of VEGFA in MDA-MB-231 cells as a proxy for miR-16-5p inhibition efficacy by a targeting TuD in the Decoy-seq configuration (CROP-seq) or commercially available TuD as compared to a non-targeting (NT) control. **g,** Dotplot showing the normalized single-cell expression of VEGFA in cells expressing a TuD targeting miR-16-5p or a non-targeting control in the Decoy-seq miRNA screen. P-values calculated using one-tailed Wilcoxon rank-sum test with mean and 99% confidence interval shown.

**Extended Fig 2.**
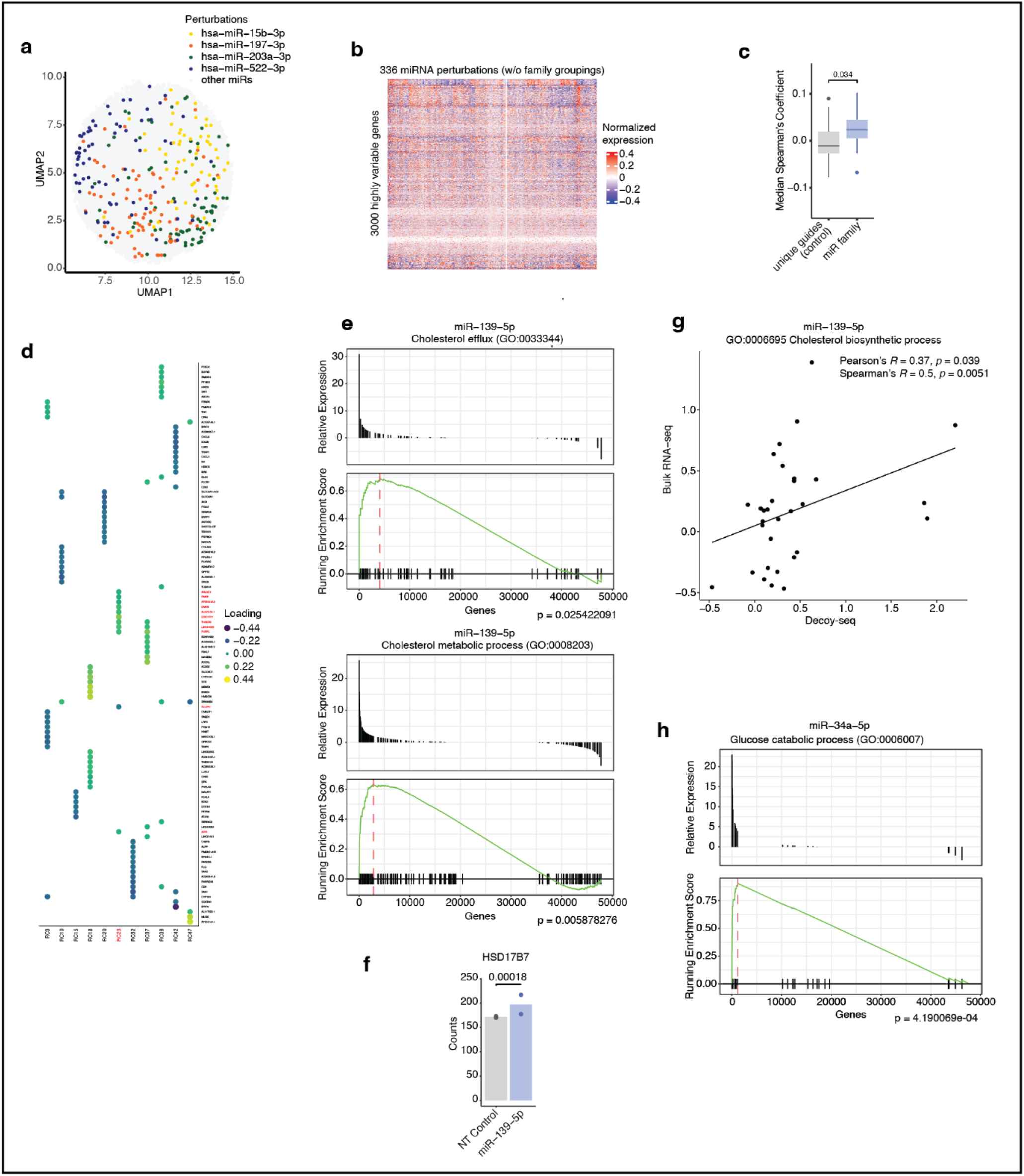
Transcriptomic phenotypes associated with miRNA perturbation. **a,** UMAP of 3000 highly variable genes across single cells illustrating that miRNA perturbations created distinct transcriptomic cell states across the transcriptome. **b,** Heatmap of normalized expression levels of 3000 highly variable genes (rows) across aggregated pseudo-bulk profiles of each miRNA perturbation (column) without family grouping (from Seurat’s SCTransform function). **c,** Median Spearman’s correlation coefficient of Decoy-seq pseudo-bulk expression profiles of perturbation of miRNAs with a shared (family) seed sequence (i.e. within a family) or equivalently sized pseudo-family made from randomly selected miRNAs with unique seed sequence. P-value calculated using Wilcoxon signed-ranked test. **d,** Rotated component loading values plotted for high loading genes (absolute value > 0.1) of RCs with at least one miRNA perturbation with significant shift across components. These genes were used in the Enrichr algorithm to identify relevant pathways. High loading genes for RC23, associated with miR-34a-5p, are highlighted in red. **e,** GSEA analysis of differentially expressed genes of the GO cholesterol efflux and metabolic process genesets shown between miR-139-5p inhibition and control. Plotted is the relative expression or the running enrichment score of all sequenced genes ordered by upregulated to downregulated via the DESeq2 Wald test statistic. P-value calculated by odds of peak enrichment score out of GSEA simulations done with random gene order, adjusted for all geneset comparisons by Benjamini-Hochberg procedure. **f,** Barplot of normalized gene expression count in a key gene in cholesterol biosynthesis, HSD17B7, from the bulk RNA-seq data of MDA-MB-231 cells transduced with a non-targeting (NT) control TuD or a TuD targeting miR-139-5p. P-values calculated in DESeq2 using the two-tailed Wald test and adjusted with the Benjamini-Hochberg procedure. **g,** Scatter plot showing Pearson’s and Spearman’s correlations between Decoy-seq and bulk RNA-seq expression profiles of miR-139-5p perturbation. The x and y-axis represent log2 fold change of genes in the cholesterol biosynthesis process GO gene-set between the miR-139-5p inhibition and NT control. Pearson’s and Spearman’s correlation coefficients R and p-values for plotted line fit (calculated using two-sided Student’s t test) are indicated. **h,** GSEA analysis of differentially expressed genes of the GO glucose catabolic process geneset shown between miR-34a-5p inhibition and control.

**Extended Fig 3.**
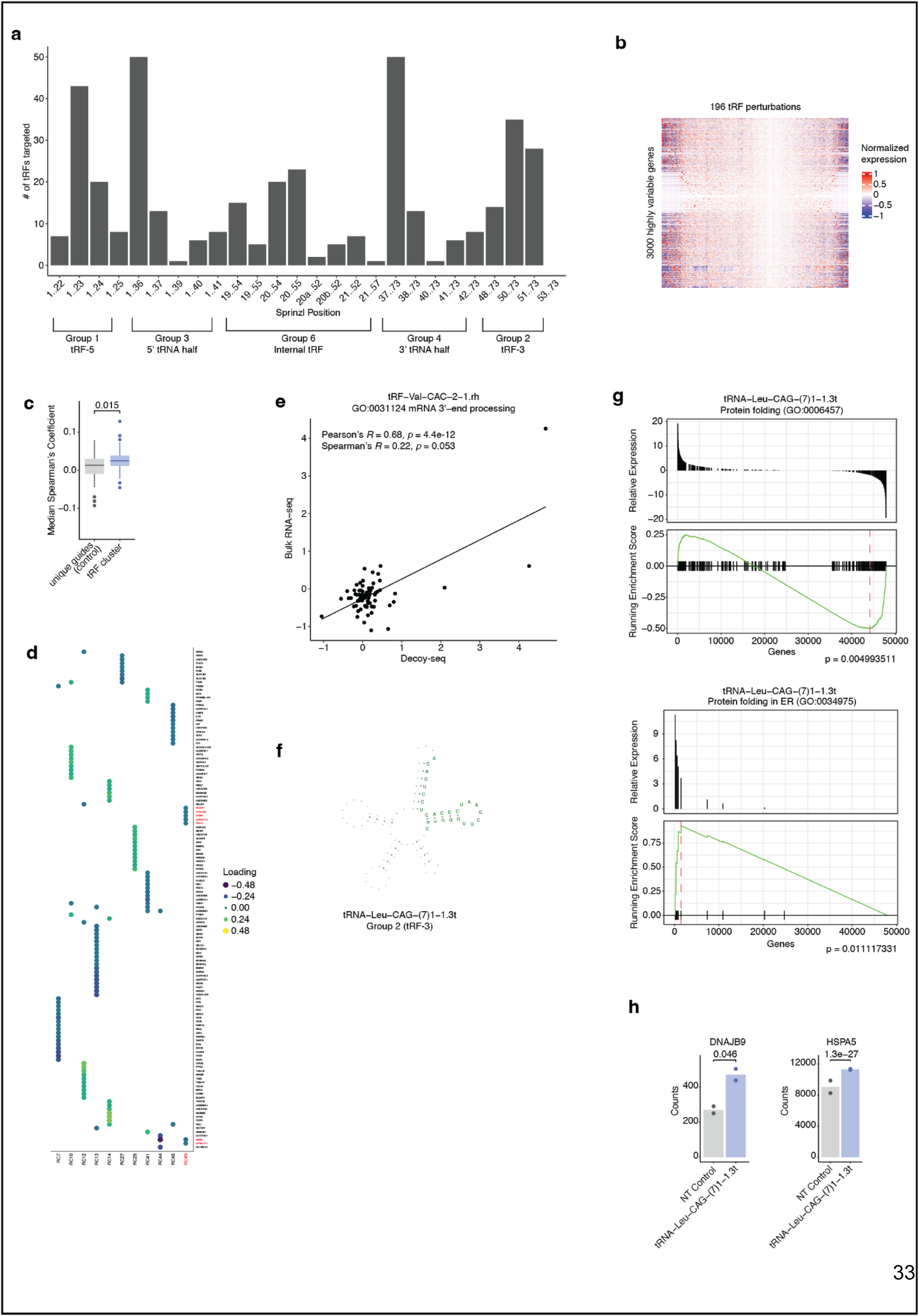
Transcriptomic phenotypes and biological pathways associated with tRF perturbation. **a,** Histogram showing the distribution of the subgroups of tRFs targeted in the Decoy-seq discovery screen. The x-axis represents the Sprinzl position (ie the base pair positions of the tRNA from which the tRF is derived). tRFs are subgrouped based on Su et. al. (2020) in ref 19. **b,** Heatmap of normalized expression levels of 3000 highly variable genes (rows) across aggregated pseudo-bulk profiles of each tRF perturbation (column) without clustering (from Seurat’s SCTransform function). **c,** Median Spearman’s correlation coefficient for Decoy-seq pseudo-bulk expression profiles of perturbation of tRFs within a cluster or equivalently sized pseudo-cluster made from randomly selected tRFs with unique sequence. P-value calculated using Wilcoxon signed-ranked test. **d,** Rotated component loading values plotted for high loading genes (absolute value > 0.1) of RCs with at least one tRF perturbation with significant shift across components. These genes were used in the Enrichr algorithm to identify relevant pathways. High loading genes for RC49, associated with the perturbation of tRNA-Val-CAC-2-1.rh, are highlighted in red. **e,** Scatter plot showing Pearson’s and Spearman’s correlations between Decoy-seq and bulk RNA-seq expression profiles of tRNA-Val-CAC-2-1.rh perturbation. The x and y-axis represent log2 fold change of genes in the GO mRNA 3’-end processing geneset between the miR-139-5p inhibition and NT control. Pearson’s and Spearman’s correlation coefficients R and p-values for plotted line fit (calculated using two-sided Student’s t test) are indicated. **f,** Illustration of the sequence position highlighted in green of tRNA−Leu−CAG−(7)1−1.3t and its subgrouping, indicating its origin within the tRNA from which it was derived. **h,** (left and middle panels) GSEA analysis of differentially expressed genes of the GO “protein folding” genesets shown between tRNA-Leu-CAG-(7)1-1.3t inhibition and control. Plotted is the relative expression or the running enrichment score of all sequenced genes ordered by upregulated to downregulated via the DESeq2 Wald test statistic. P-value calculated by odds of peak enrichment score out of GSEA simulations done with random gene order, adjusted for all geneset comparisons by Benjamini-Hochberg procedure. (right panel) Barplots of normalized gene expression count in two key regulator genes in IRE1-mediated unfolded protein response, DNAJB9 and HSPA5, from the bulk RNA-seq data of MDA-MB-231 cells transduced with a non-targeting (NT) control TuD or a TuD targeting tRNA-Leu-CAG-(7)1-1.3t. P-values calculated in DESeq2 using the two-tailed Wald test and adjusted with the Benjamini-Hochberg procedure.

**Extended Fig 4.**
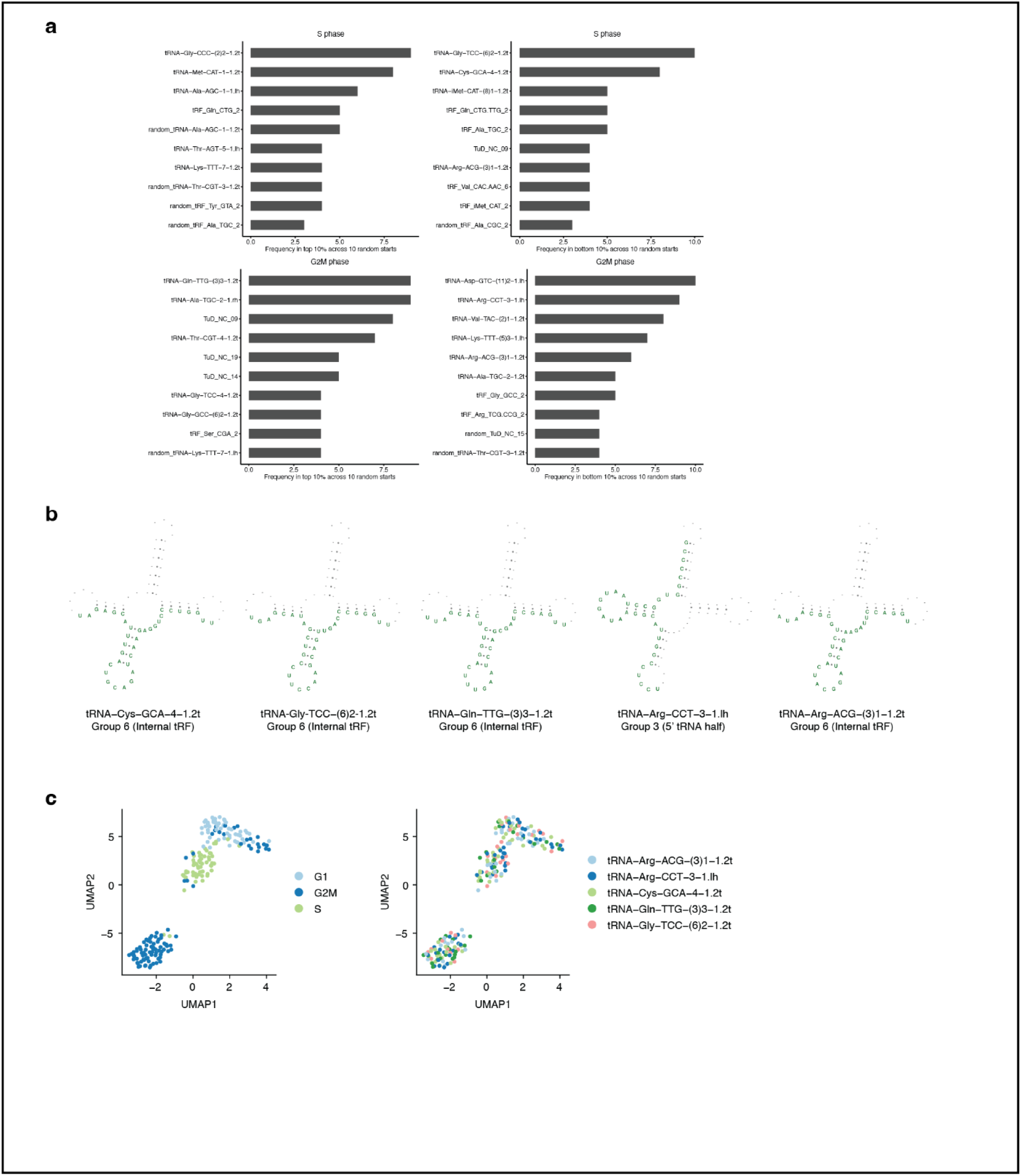
Changes in cell cycle progression in relation to tRF perturbations. **a,** Frequency of tRF perturbation that ranked within the top or bottom 10% based on their median cell cycle scores in the S or G2/M phases in the Decoy-seq screen data. The top two tRFs from these four plots shown were selected for further testing in Fig 4. Only the validated tRFs were shown in Fig 4. **b,** Illustration of the sequence position highlighted in green of each tRF validated in Fig 4 and their subgrouping, indicating their origin within the tRNA from which they were derived. **c,** UMAP analysis depicting the cell cycle state of each cell based on the expression of cell cycle genes in the Decoy-seq data (left), along with the assignment of tRF perturbations (right) selected for further experimental validation in Fig. 4.

